# An Integrated Computational Approach to Predict and Characterize Emerging Mutations in the Japanese Encephalitis Virus Envelope Protein

**DOI:** 10.64898/2026.05.26.727781

**Authors:** Hariprasad Thippeswamy, Kuralayanapalya Puttahonnappa Suresh, Rajan Kumar Pandey, Yamini Sri Sekar, Varsha Ramesh, Navnath Kamble, Azhahianambi Palavesam, Sharanagouda S. Patil, Jagadish Hiremath

## Abstract

Japanese encephalitis virus (JEV) causes significant encephalitis across the Asia-Pacific region. Current vaccines target historical genotype III strains, but emerging genotypes,potentially driven by vaccine-mediated selective pressure, threaten vaccine effectiveness through altered envelope protein sequences that may reduce antibody cross-neutralisation. This study employed integrated sequence and structural analyses to identify E protein mutations affecting neutralising antibody binding and protein stability. The study curated JEV polyprotein sequences from NCBI, performed multiple sequence alignment, and used Shannon entropy to pinpoint highly variable positions. Mutations occurring at ≥1% frequency within high-entropy regions were selected for analysis. From 34 initially identified mutations, four candidates were prioritized based on structural stabilization potential. Mutations were evaluated through FoldX stability predictions, molecular docking with antibody 2H4 using HADDOCK3, and molecular dynamics simulations. Binding energies were calculated using MM-GBSA analysis. Results demonstrated that all mutant E-2H4 complexes remained stable during simulations, with root-mean-square deviation plateauing after equilibration and minimal localized changes in root-mean-square fluctuation. These findings suggest that EDIII substitutions represent important candidates for further investigation to understand genotype-specific variations and inform next-generation vaccine development strategies against emerging JEV strains.

## Introduction

Japanese encephalitis virus (JEV) causes Japanese encephalitis (JE) in humans and animals (Oliveira et al., 2018). JEV belongs to the genus Flavivirus, and this vector-borne disease is mainly transmitted through mosquito bites (Solomon et al., 2002). Although most human JEV infections are asymptomatic, about 1% of cases present with clinical symptoms. JE can have a death rate of up to 18%, and in about 45% of non-fatal cases, long-term neurological or psychological problems may occur (Weng et al., 2018). Japanese encephalitis prevalence is rising due to climate change and intensive irrigation creating vast mosquito breeding grounds, alongside the expansion of pig farming, which provides a massive viral reservoir. These environmental shifts are compounded by improved diagnostic reporting and gaps in vaccination coverage, allowing the virus to spread into previously unaffected geographical regions (Erlanger et al., 2009). However, there is no proven treatment for this disease, despite its rising incidence and threat to human health. This situation urges to increase our understanding of JEV, including its genetic structure, pathophysiology, diagnostic methods, vaccine development, and potential antiviral strategies, to better prepare for future outbreaks (Li et al., 2025). Vaccination remains essential for prevention. All JE vaccines approved for commercial use (JE-VAX, Ixiaro, Imojev, SA 14-14-2) are based on the Genotype III (GIII) strains (Yun et al., 2014). JE-VAX can produce neutralising antibodies against JE genotypes I through IV (Erra et al., 2013), but cannot effectively induce neutralising antibodies against both Genotype V (GV) Muar and GV XZ0934, a JEV isolate from a Chinese mosquito pool (Honjo et al., 2019; Cao et al., 2016). Compared to other JEV genotypes, the GI-based vaccine generates fewer cross-neutralising antibodies against Genotype V Muar (Tajima et al., 2021). To effectively prevent GV JEV infection, a vaccine specifically targeting GV JEV must be developed.

JEV is an enveloped positive-sense single-stranded RNA virus with about an 11-kb genome capped at the 5′ end but lacking a 3′ poly-A tail. Its single open reading frame encodes a 3400 amino acid polyprotein that is cleaved into structural proteins (C, prM/M, E) and seven non-structural proteins (NS1–NS5) (Yun et al., 2009). The E(Envelope) protein is the main viral attachment protein and fusion machinery. In JEV, the distal tip of domain III (DIII) is believed to engage host receptors that mediate host-cell entry (Luca et al., 2011).

The E protein folds into three distinct ectodomains (Domains I–III, or DI–DIII), along with a C-terminal stem and a transmembrane (TM) region (Ren et al., 2007; Ishida et al., 2023). Domain I (DI) forms a central beta-barrel, while domain II (DII) appears as an elongated “finger” with the conserved fusion loop at its tip. Domain III (DIII) resembles an immunoglobulin-like module projecting from the virion surface (Kumar et al., 2022). The JEV E protein has a conserved N-linked glycosylation site at Asn-154 for receptor attachment and forms head-to-tail homodimers with a compact DIII–DI interface; nearby conserved histidines may facilitate pH-dependent fusion (Huang et al., 2024). Mutations in these receptor-binding regions can drive antibody evasion and inform therapeutic or vaccine design. Antibody–antigen interactions are a crucial component of the immune response to pathogens, and understanding their structural basis can aid in the design of more effective therapeutics (Boyden, 1966). JEV can be strongly neutralised by monoclonal antibodies 2F2 and 2H4. These provide a high-affinity template for humanisation, as their binding regions (CDRs) can be grafted onto human frameworks to bypass the HAMA response. Their mature hybridoma technology enables rapid development of antibodies with exquisite specificity, making them indispensable for studying cross-reactivity between JEV and ZIKV. While unsuitable for direct human therapy, they serve as reliable, cost-effective tools for diagnostic assays where foreign protein immunogenicity is not a concern (Liu et al.,2025). Structural studies show that they target unique quaternary epitopes across multiple envelope proteins, offering strong therapeutic potential (Qiu et al., 2018). Mutations in the functional domain of antigen-binding interfaces can significantly change binding strength. Studying these changes through antibody–antigen docking helps in identifying key amino acids essential for immune escape. Such insights guide the redesign of protein interfaces for therapeutic purposes (Studer et al., 2013). Following docking, Molecular Dynamics (MD) simulations and MM-GBSA analyses of an antigen-antibody complex are essential for refining computationally predicted docked structures and providing a dynamic, atomic-level view of the binding process. While molecular docking predicts static poses, MD simulation and MM-GBSA offer insights into flexibility, binding stability, and energetics under physiological conditions, which are vital for accurate analysis and therapeutic design (De et al., 2016). This comprehensive research study emphasises that understanding patterns of mutation accumulation is essential for predicting potentially harmful mutations, as these patterns have not yet been explored in the envelope protein of JEV. It involves addressing two key questions: (a) which types of mutations are more likely to occur, and (b) which residue positions or regions are more prone to mutation? To answer these, this study seeks to systematically screen and validate mutations within the JEV envelope protein, identifying critical molecular targets for the development of pan-genotypic vaccines and therapeutics.

## Methods

### 2.1.1 Envelope Protein Retrieval and Processing

Complete polyprotein sequences of diverse Japanese Encephalitis Virus (JEV) strains were retrieved from the NCBI Virus repository (https://www.ncbi.nlm.nih.gov/labs/virus/vssi/#/) following a comprehensive literature-guided strain selection strategy. To ensure dataset quality and eliminate redundancy, sequences were filtered to remove duplicates, truncated or partial entries, non-structural elements irrelevant to the envelope protein, and the membrane (M) protein region. A command-line BLASTp search was then performed using the reference envelope protein sequence (RefSeq ID: NP_775666.1) as a query against the filtered dataset. The envelope region was computationally extracted from each retained strain using homology to the reference sequence. To maintain high dataset specificity, we explicitly excluded phylogenetically closely related flaviviruses such as Murray Valley encephalitis virus (MVEV), West Nile virus (WNV), and other non-JEV members of the JEV serocomplex

#### 2.1.2 Structural Template Identification and Homology Modeling

To establish a high-fidelity spatial coordinate system and precisely define the 3D scaffold for mutational analysis, the reference envelope protein was modelled using MODELLER v10.4 (Sali and Blundell, 1993). Multiple homologous flaviviral E protein crystal structures (PDB IDs: 3P54, 5MV1, 5MV2) were selected as templates to enhance model reliability through multi-template alignment and spatial restraint optimisation. Template selection was based on high sequence similarity and structural conservation within the Flavivirus genus. The resulting 3D model was validated using Ramachandran plot analysis to assess the quality of backbone dihedral angles, ensuring stereochemical reliability. This validated model was then used as a reference scaffold for *in silico* mutagenesis at selected residues, providing a structurally consistent foundation for subsequent computational characterisation of mutation-induced effects.

### 2. 2 Sequence Examination

#### 2.2.1 Sequence Pre-processing

Extracted envelope protein sequences were subjected to rigorous pre-processing to ensure analytical consistency and eliminate artefacts. Sequences containing ambiguous amino acid codes (e.g., X, B, Z) were excluded to prevent misalignment or erroneous mutation calls during downstream analyses. To maintain structural comparability with the reference sequence, entries were filtered by amino acid length (500 residues), retaining only those within a ±5-residue window relative to the reference envelope protein. Subsequently, redundant sequences were clustered, and identical entries were collapsed to a single representative per cluster. This deduplication step reduced computational complexity and ensured that the resulting dataset accurately captured the unique sequence diversity required for mutation frequency and entropy analyses

#### 2.2.2 Identification of Position-Specific Mutation Count

To quantify sequence variability across envelope protein, a high-resolution positional analysis was conducted to map the distribution of substitutions in the scaffold. Multiple sequence alignment was conducted using the MAFFT61 algorithm (Katoh and Standley, 2013) with its default settings, with its Fast Fourier Transform (FFT) to rapidly identify homologous regions, followed by a progressive alignment guided by a similarity tree and optional iterative refinement to maximize accuracy. The aligned output was converted into a structured data frame, with the first column listing the viral strain and the subsequent columns recording the amino acid at each residue position across the sequence. When clustering was applied before alignment, an additional column was included to indicate the size of each cluster. Positions corresponding to gaps (“-”) in the reference sequence were excluded, as these represent insertion events. Each residue position in the alignment was then compared with the reference sequence to identify mutations. Mutation counts per site were tallied, and mutation frequencies were calculated as the ratio of the total mutation count to the number of sequences analyzed. For cluster-based alignments, the mutation count from the alignment was weighted by multiplying it by the associated cluster size. The weighted total was then divided by the overall number of sequences in the dataset to obtain the actual mutation frequency.

#### 2.2.3 Detection of high-frequency mutations

To evaluate sequence variability, Shannon entropy (SE) was calculated for each residue position in the alignment. SE quantifies the degree of uncertainty or randomness within a dataset and is defined by the formula (Shannon, 1948).

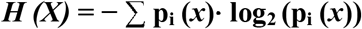

where *H* (*X*) denotes the entropy at a given alignment position *X*, and *p*_*i*_ (*x*) represents the probability of observing residue *x* at that position. The summation is performed over all residues at that site. Positions with high entropy values reflect greater variability, whereas low entropy values indicate conserved sites. The distribution of SE values was examined through a density plot, and the noticeable drop in density was selected as the threshold to distinguish low- and high-entropy positions. Residues falling into the high-entropy category were further analyzed, and mutations occurring with a frequency of ≥1% (Sharma et al., 2024) within these positions were classified as high-frequency mutations.

### 2.3 Computational Framework for Predicting the Structural and Immunological Impact of Envelope Protein Mutations

Evaluating the structural and immunological perturbations induced by substitutions in the Japanese Encephalitis Virus (JEV) envelope (E) protein requires a multi-scale computational framework to quantify mechanistic shifts in protein stability and antibody-binding affinity. The implemented pipeline integrates mutation-specific thermodynamic stability profiling, molecular docking, and atomistic molecular dynamics (MD) simulations to create a high-fidelity map of the protein’s functional landscape. The selected mutations were identified according to sequence-based screening and supported by literature as functionally relevant alterations within the receptor-binding domain (RBD). These mutations may be implicated in modulating host receptor interaction and facilitating immune evasion, thereby influencing fold stability.

#### 2.3.1 Structural Stability Prediction via FoldX

Quantifying the thermodynamic consequences of amino acid substitutions requires a high-resolution force field to evaluate the localized energetic shifts within the folded domain. This is critical as even minor structural perturbations can significantly alter the protein’s folding kinetics and proteolytic stability, factors that cannot be deduced from primary sequence data alone. Building upon this requirement, the FoldX 5.0(Schymkowitz et al., 2005) empirical force field was utilized to calculate the change in the Gibbs free energy of folding (ΔΔG) between the wild-type and mutant states. FoldX estimates ΔΔG using an empirical force field that incorporates van der Waals interactions, hydrogen bonding, electrostatics, solvation effects, and entropic contributions. As per the scale of the ΔΔG values, mutations were classified as stabilizing (ΔΔG < 0), destabilizing (ΔΔG > 0), or neutral (ΔΔG ≈ 0). Each mutant structure was prepared using the FoldX “BuildModel” module, followed by hydrogen bond network optimization and restrained energy minimization to ensure structural realism before downstream molecular dynamics simulations. **-**

#### 2.3.2 Evaluation of Mutational Impact on Antigen-Antibody Binding via HADDOCK3

To investigate how each mutation influences antigen–antibody interaction, antigen–antibody docking was performed using the HADDOCK3 platform (Ambrosetti et al., 2020). The process began with topology generation (topoaa), followed by rigid-body docking (rigidbody) using ambiguous interaction restraints (AIRs) according to ProABC-2-predicted paratope and NMR-defined epitope regions. CAPRI-based scoring (caprieval) was applied iteratively throughout the workflow to evaluate docking quality via metrics such as iRMSD, Fnat, and DockQ. Semi-flexible refinement (flexref) and energy minimization (emref) further optimized the interface geometry. Clustering was performed using the fraction of common contacts (clustfcc), and the top models from each cluster were selected (seletopclusts) for detailed analysis. Contact mapping (contactmap) was used to visualize interfacial interactions. Restraint definitions were rigorously validated to ensure compatibility, and top-ranked docked complexes were advanced to molecular dynamics simulations for final stability and binding energy assessments. The E protein served as the antigen, while the neutralizing Fab fragment of monoclonal antibody 2H4 (PDB ID: 5YWF) was used as the antibody component. Each mutant variant was modeled individually onto the validated E protein structure, and all docking input files were prepared in accordance with HADDOCK3 preprocessing guidelines: heteroatoms were removed, alternate conformations were resolved, chain IDs were uniquely assigned, and non-overlapping residue numbering was enforced. As HADDOCK3 is employed to capture both structural accuracy and interaction fidelity between mutant JEV envelope proteins and the 2H4 neutralizing antibody.

#### 2.3.3 Molecular Dynamics Simulations

To evaluate the conformational stability and interaction fidelity of antigen–antibody complexes, molecular dynamics (MD) simulations were performed using GROMACS v2025.3. System preparation followed the standard GROMACS workflow: topology generation (pdb2gmx), box definition (editconf), solvation (solvate), ion addition (genion), minimization (grompp and mdrun), and equilibration phases. Post-docking, the resulting antigen–antibody complexes were solvated, neutralized with 0.15 M NaCl, and subjected to energy minimization, NVT/NPT equilibration, and 100 ns production runs using the AMBER19SB force field (Tian et al.,2019). Simulations were executed with GPU acceleration for parallel processing, and trajectories were recorded for downstream analysis.

#### 2.3.4 Binding Free Energy Estimation through MM-GBSA

Binding energetics of the docked complexes were quantified using the MM-GBSA method implemented in the gmx_MMPBSA (Valdés-Tresanco., 2021) tool interfaced with GROMACS. A trajectory protocol was employed to extract antigen and antibody coordinates directly from 100 ns production trajectories. The stable frames from stable trajectory windows were used to compute ΔG_bind, applying the Generalized Born (GB) model with a 0.15 M salt concentration. Binding energy (ΔG_bind) was calculated as:

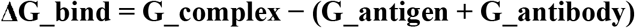

where each term includes molecular mechanics energy (EMM), solvation free energy (Gsolv), and non-polar contributions.

## Results

### 3.1 Sequence Examination

Among the 1,210 input sequences, 45 (3.7%) were removed due to ambiguous residues, and 9 (0.7%) failed the length filter, leaving 1,156 usable sequences. After deduplication/clustering, 370 unique sequences (370 clusters) remained; the largest cluster comprised 320 sequences. We then performed multiple sequence alignment of the 370 unique sequences together with the reference. Gaps at reference insertions are removed, and per-position mutations and frequencies are computed from the aligned residues. Across the cohort (n = 371 sequences), 2,994 amino-acid changes were detected spanning 313/500 positions (62.6%).

#### 3.1.1 Identification of high-frequency mutations

Applying a ≥5% frequency screen identified 34 recurrent “hotspot” positions, with prominent peaks at 169, 327, 366, 222, and 129 (S1) that identify the drivers of evolution. Per-position Shannon entropy (H) was calculated across the 500 sites to quantify residue-level diversity. Using the sharp decline in H density **(Figure 2)** as a data-driven cutoff (H* = 0.0271), 196/500 (39.2%) positions were labeled “high-entropy.” Mutations occurring at high-entropy sites with allele frequency ≥1% were classified as high-frequency mutations that contribute to the diversity of the population. This filter yielded 107 high-frequency mutations mapping to 92 positions. The most enriched single-site substitutions included 366 A→S, and 129 T→M, 327 S→ Q, 138E→K as literature based mutant. The entropy confirms a right-skewed landscape with most residues near H≈0 and a tail of highly variable sites

**Figure 1.**
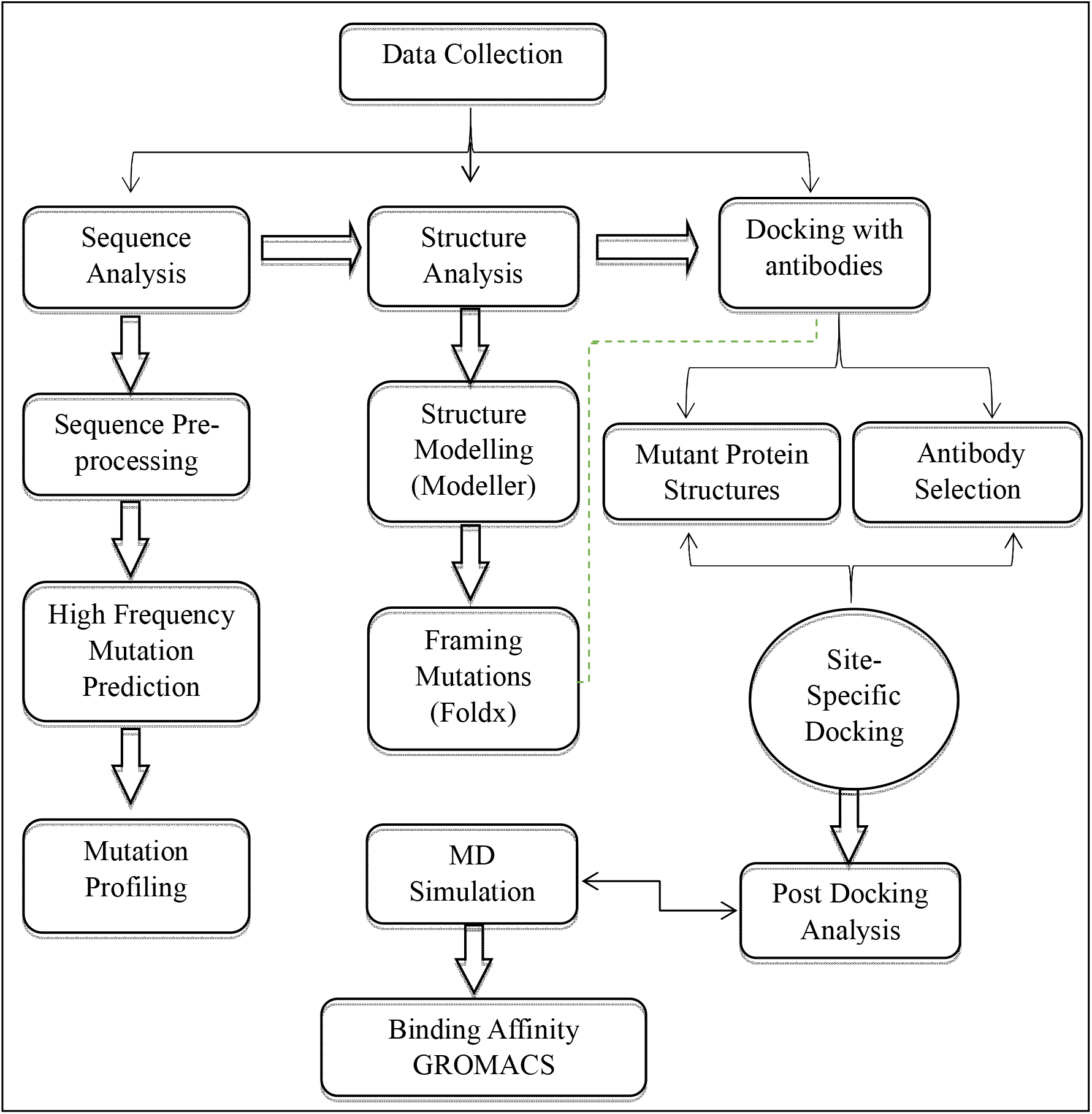
An illustration of the comprehensive framework employed in this study for systematic evaluation of mutation-driven effects on JEV E protein structure and antibody interaction.

**Figure 2.**
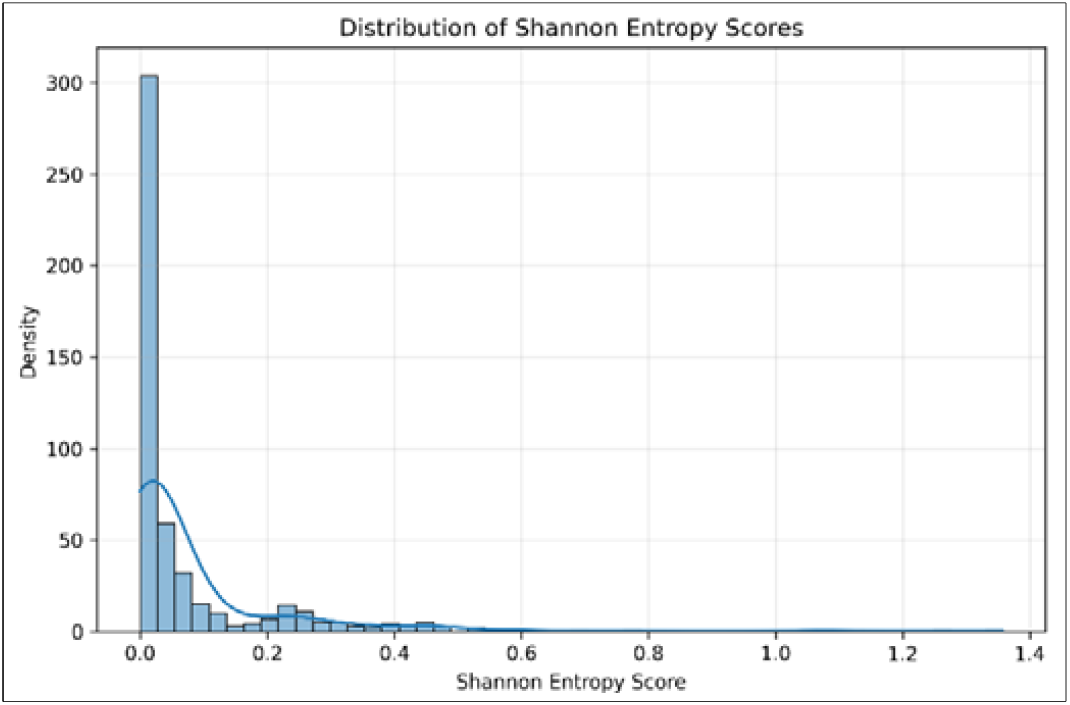
This figure illustrates the distribution of per-residue Shannon entropy (H) across the envelope protein of JEV, highlighting regions of elevated sequence variability.

### 3.2 Structure Modeling and Computational Mutagenesis of JEV Envelope Protein

The Japanese encephalitis virus envelope (E) protein structure was modeled based on the RefSeq strain NP_775666.1 and subjected to comprehensive structural validation using Ramachandran plot analysis, which confirmed favorable backbone conformations consistent with high-quality protein structures. The 500 residue E protein is organized into three ectodomains (DI, DII, DIII), a stem region, and a transmembrane anchor. FoldX computational mutagenesis was performed on four strategically selected positions to evaluate their contribution to structural stability through calculation of free energy changes (ΔΔG).

#### 3.2.1 FoldX Computational Mutagenesis Results

The computational analysis revealed a stabilizing mutational landscape, with four mutations conferring enhanced stability (ΔΔG < 0). The mutations spanned all three envelope domains, with Domain III showing the highest concentration of stabilizing substitutions and the most favorable average ΔΔG value. Position 327: a stability hotspot; S327Q (ΔΔG = −0.97 kcal/mol) markedly stabilizes Domain III. Glutamine at 327 likely strengthens local hydrogen bonding via its longer side chain, while threonine adds a methyl group that improves hydrophobic packing while preserving the hydroxyl’s polar contacts. The fact that two chemically distinct substitutions converge on stabilization indicates a structurally permissive microenvironment at 327. Because Domain III mediates receptor engagement and hosts neutralizing epitopes, stabilizing changes here could elevate fold stability while reshaping antigenic recognition. Domain I stabilizers; E138K (ΔΔG = −0.23 kcal/mol) reverses charge and likely optimizes electrostatics within the Domain I β-barrel core. It has been observed during passage, where it restored entry and coincided with attenuation of neurovirulence. T129M (ΔΔG = −0.21 kcal/mol) places a bulkier hydrophobe near the DI–DII hinge, improving core packing and potentially filling a latent cavity. Contextually, A366S (ΔΔG = −0.15 kcal/mol) is mildly stabilizing in Domain III—consistent with formation of compensatory hydrogen bonds

#### 3.2.2 Docking performance of JEV E protein mutants

HADDOCK3 was used in the docking of the JEV envelope protein mutants. FCC clustering consistently identified a dominant top-ranked cluster **(Table 2)** for each mutant, indicating stable and reproducible antigen–antibody binding modes. Mutations introducing charge or polarity changes (E138K, S327Q, A366S) showed greater modulation of interface energetics and variability. None of the mutations abolished antibody binding, suggesting that the 2H4 antibody maintains recognition across these mutants, although with mutation-specific differences in binding strength and interface organization. Electrostatic interactions played a dominant role in variants such as E138K, consistent with the introduction of a positively charged residue, whereas mutations like T129M primarily affected hydrophobic packing. Variants S327Q exhibited increased interface flexibility, reflected in broader cluster variability, while A366S retained stable binding with only minor local rearrangements.

**Table 1.**
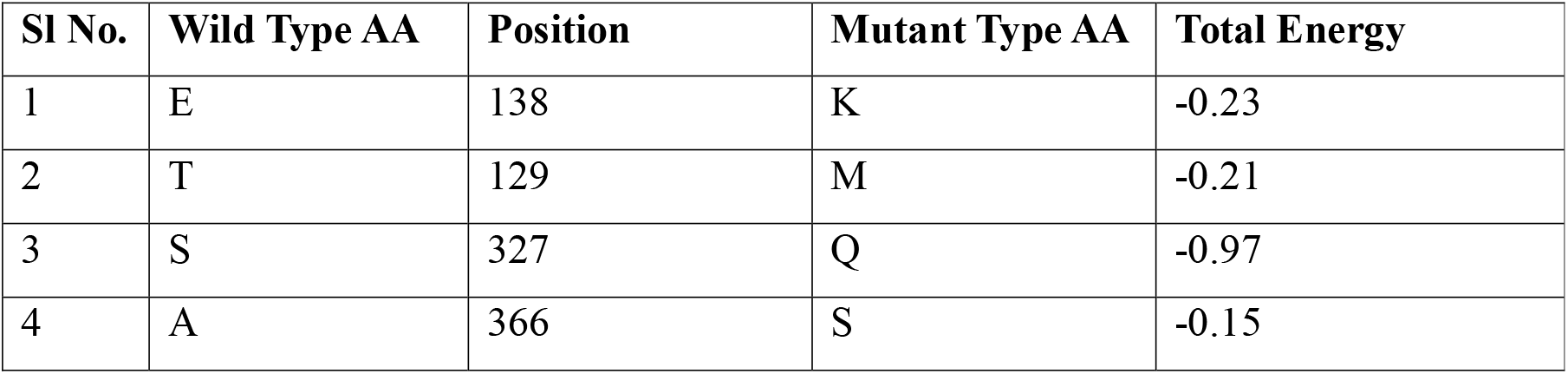
FoldX mutagenesis on JEV envelope protein sequences.

| Sl No. | Wild Type AA | Position | Mutant Type AA | Total Energy |
| --- | --- | --- | --- | --- |
| 1 | E | 138 | K | -0.23 |
| 2 | T | 129 | M | -0.21 |
| 3 | S | 327 | Q | -0.97 |
| 4 | A | 366 | S | -0.15 |

**Table 2.** HADDOCK3 docking statistics for JEV envelope protein mutants docked to the 2H4 antibody.

| Mutant | HADDOCK score | iRMSD (Å) | Fnat | BSA (Å<sup>2</sup>) | van der Waals | Electrostatics |
| --- | --- | --- | --- | --- | --- | --- |
| E138K | -62.86 | 0.951 | 0.76 | 1925 | -71.79 | -291.576 |
| A366S | -39.35 | 0.56 | 0.85 | 1886 | -37.43 | -231.86 |
| S327Q | -37.95 | 0.47 | 0.92 | 1844 | -47.75 | -143.46 |
| T129M | -30.21 | 0.71 | 0.77 | 1768 | -64.53 | -195.77 |

#### 3.2.3 Assessment of mutational stability through MD simulation

To evaluate the structural stability of the docked complexes between the 2H4 antibody and the JEV envelope antigen mutants. MD simulations indicate that all four JEV E protein mutants (T129M, E138K, S327Q, A366S) maintain stable complexes with the 2H4 neutralizing antibody. Backbone RMSD rose during early equilibration (∼10–20 ns) and then plateaued for every system, with Cα RMSD generally staying within ∼2–3 Å of the starting docked structure **(Figure 3)**. A366S showed the smallest fluctuations (stabilizing near ∼2 Å), while E138K displayed a slightly larger initial shift (∼3 Å) before stabilizing, consistent with local adjustment to the charge-reversal substitution. T129M and S327Q reached similarly steady RMSD profiles, indicating no major structural drift or loss of complex integrity.

**Figure 3.**
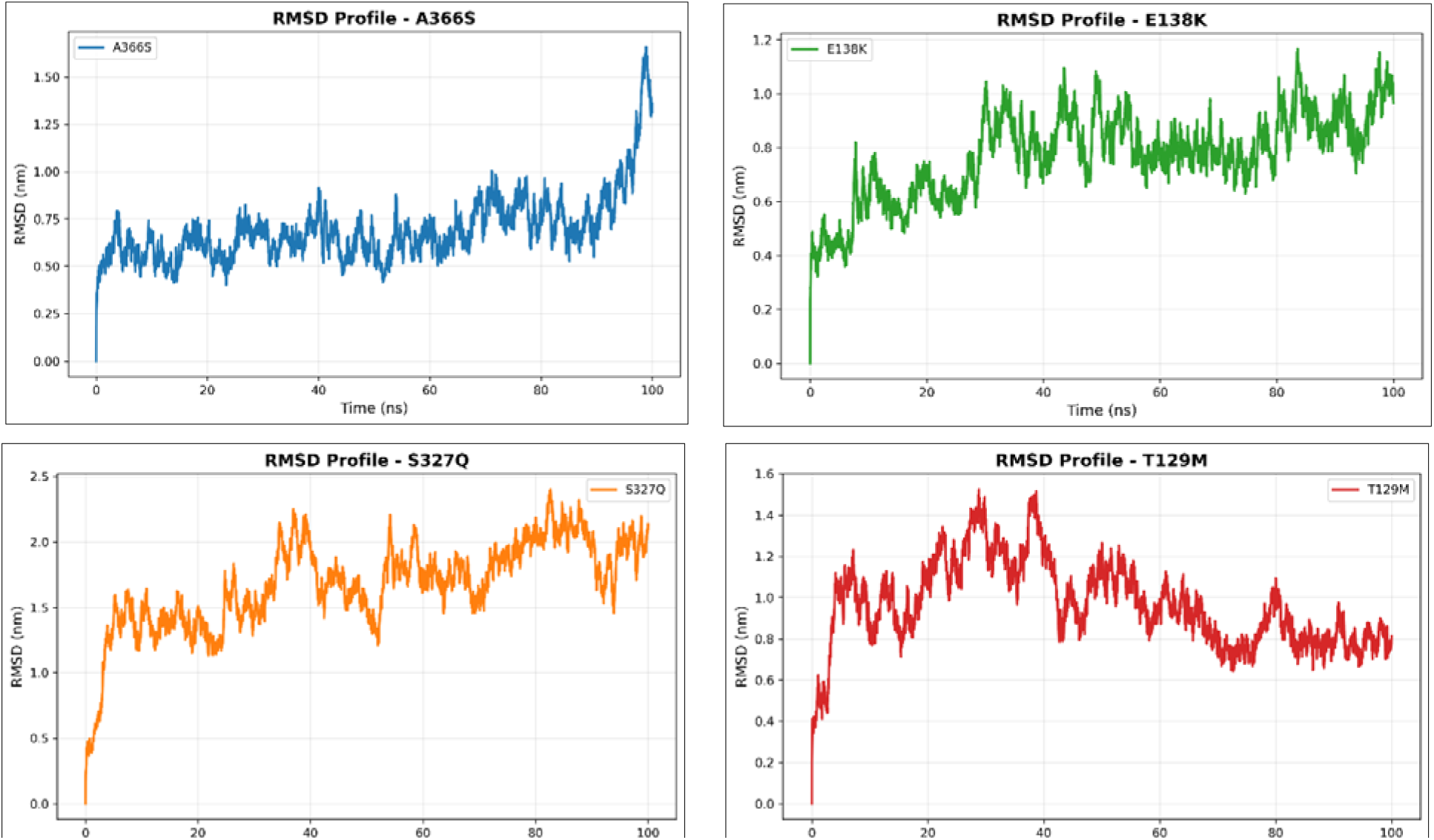
RMSD Profiles of 2H4 Antibody–JEV E Protein Mutant Complexes S327Q exhibited the highest structural deviation (>2.0 nm), while A366S showed late-stage stabilization; T129M and E138K reached equilibrium with stable binding over 100 ns MD simulations.

RMSF profiles **(Figure 4)** were broadly similar across mutants, with higher mobility confined to exposed loops and domain junctions and a relatively rigid core within domains I and III. Mutation effects were localized: S327Q produced a modest increase in flexibility around residue 327, and A366S caused a subtle increase in a nearby domain III loop. In contrast, the domain II substitutions (T129M, E138K) had minimal impact beyond their immediate sites, with RMSF curves largely overlapping across the rest of the protein. The antibody remained stable overall, with expected higher fluctuations mainly in selected CDR loops.

**Figure 4.**
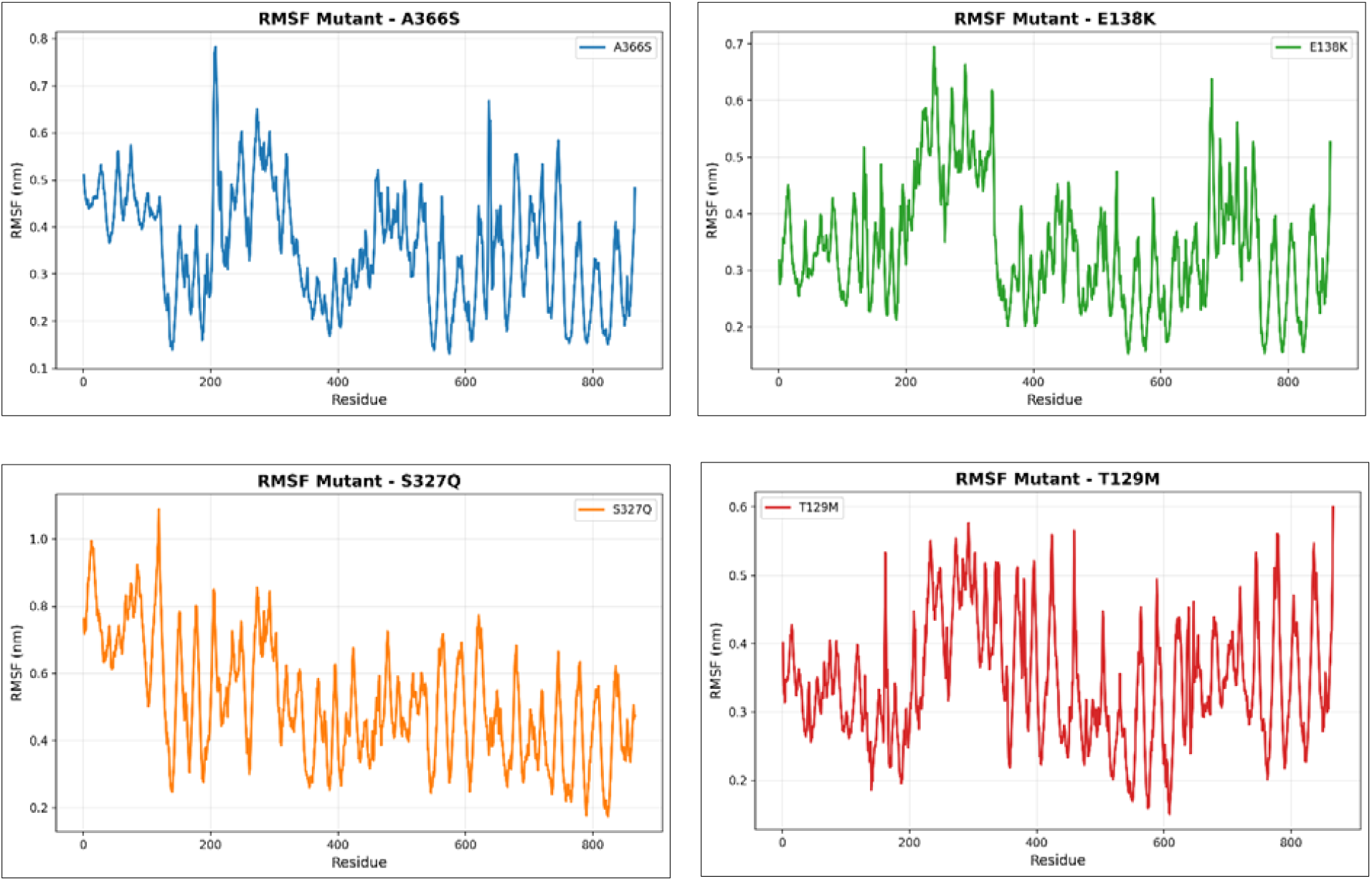
RMSF Profiles of 2H4 Antibody–JEV E Protein Mutant Complexes revealed elevated flexibility in the S327Q and A366S complexes, particularly at interface loops, while T129M and E138K maintained lower fluctuations, indicating more stable antibody–antigen interactions.

**Figure 5.**
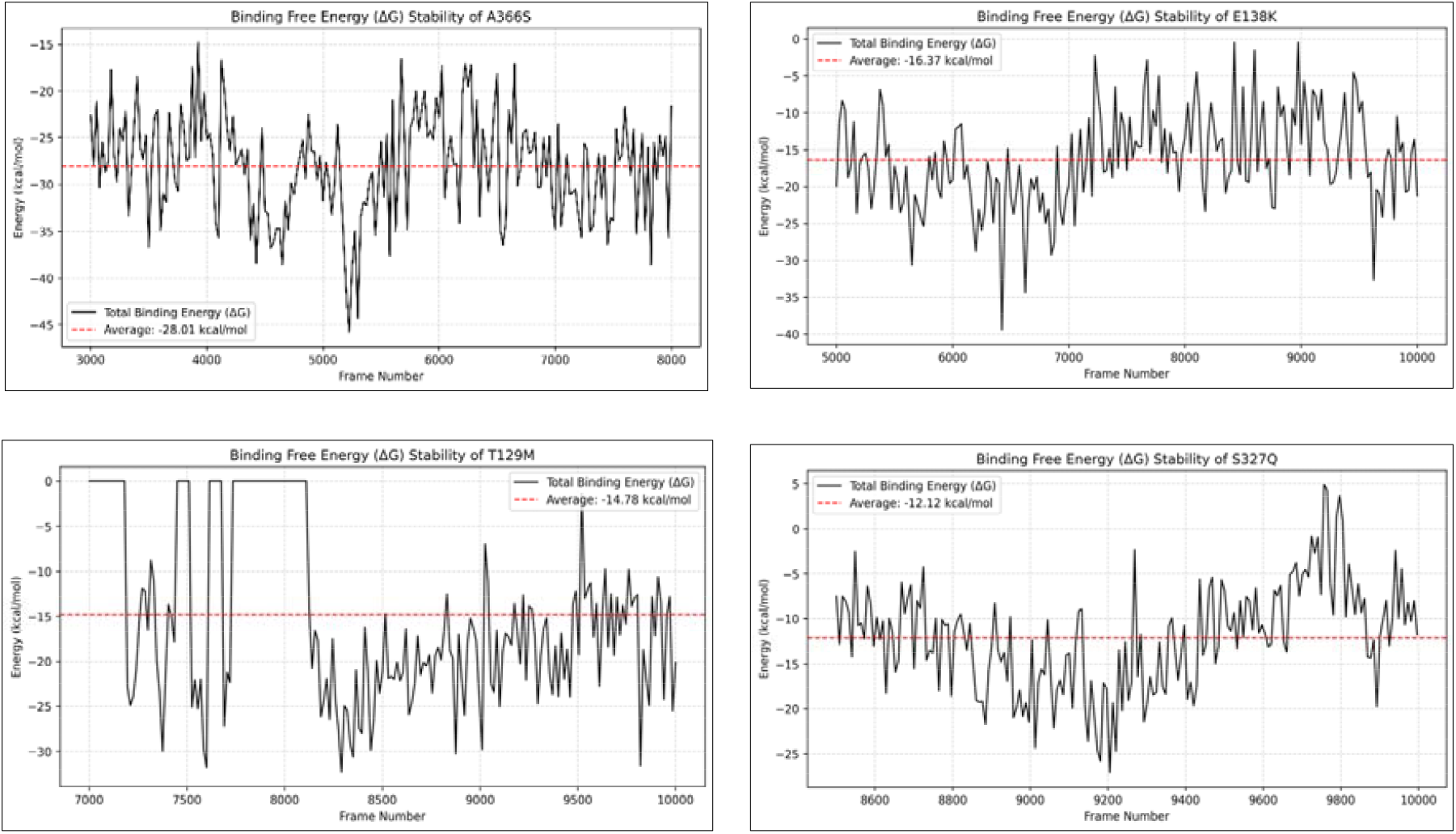
Binding Free Energy (ΔG) Profiles of Docked Mutant Complexes showed the most favorable binding energy for T129M, E138K, followed by A366S, and while S327Q exhibited the weakest binding, reflecting mutation-specific effects on antigen–antibody affinity.

MM-GBSA binding free-energy analysis showed favorable negative ΔG_bind values for all mutant–2H4 complexes, confirming stable ligand binding. A366S exhibited the strongest binding stability, with the most negative average ΔG_bind of −28.01 kcal/mol. E138K showed moderate stability with an average of −16.37 kcal/mol, while T129M displayed comparable intermediate binding energy of −14.78 kcal/mol, despite early fluctuations. S327Q had the weakest binding profile, with the least negative average ΔG_bind of −12.12 kcal/mol and greater late-stage variation. Overall, no complex showed progressive loss of binding, supporting stable 2H4 association across all mutants with mutation-specific energetic differences

## Discussion

Japanese encephalitis remains a major pediatric cause of viral encephalitis across the Asia– Pacific, and because there is no specific antiviral therapy, prevention depends largely on vaccination and vector control. A growing challenge is that licensed JE vaccines are derived from historical genotype III (GIII) strains, whereas genotype I (GI) and other non-GIII lineages (including IV/V) have expanded in circulation; sequence divergence in the envelope (E) protein—the main target of neutralizing antibodies, can shift antigenicity and has been linked to reduced neutralization of non-GIII viruses by vaccine-induced sera. In the first of its kind in the JEV, this study examines genotype-representative substitutions with potential antigenic consequences. We analyzed four E protein mutations (T129M, E138K, S327Q, A366S) using FoldX stability calculations, docking, molecular dynamics (MD), and MM-GBSA binding free energy estimation. These highlights show that each site may influence structural integrity and antibody engagement.

Across methods, a consistent pattern emerged: domain III substitutions (S327Q and A366S) produced the most evident local dynamical changes and the clearest reductions in predicted antibody binding, whereas T129M (domain II) and E138K (domain I) were comparatively conservative with respect to overall fold stability and antigen recognition. In MD, the EDIII variants increased flexibility in their immediate neighbourhoods, S327Q around the loop encompassing residue 327 and A366S near the C-terminal/366-adjacent segment, suggesting local loosening rather than global stabilization; this is important because modest reweighting of conformations at exposed epitope surfaces can weaken antibody complementarity without disrupting the broader E scaffold (Wei et al., 2019). This structural signal was mirrored by docking and MM-GBSA, where both EDIII variants showed less favourable binding relative to wild-type, with ΔG bind shifting by a few kcal/mol (typically ∼2–4 kcal/mol) toward weaker affinity, consistent with loss or rearrangement of key interface contacts. FoldX further supported a mild destabilizing tendency for these EDIII mutations (positive ΔΔG_folding on the order of ∼0.5–1 kcal/mol), which is compatible with slightly increased conformational plasticity and provides a plausible mechanism for partial immune escape driven by subtle epitope reshaping (Xu et al., 2023).

By contrast, the domain II mutation T129M exhibited minimal structural or energetic deviation from wild-type across the same analyses. Its RMSD and RMSF differences are consistent with this stability, docking and MM-GBSA predicted negligible impact on antibody binding (often within typical uncertainty, e.g., <∼0.5 kcal/mol difference), and FoldX estimated a near-neutral effect on folding stability (ΔΔG_folding ∼0). Together, these results suggest that T129M alone is unlikely to be a primary driver of antibody escape in this binding model, even if it may influence other phenotypes (e.g., aspects of fusion dynamics or host adaptation) that are not directly captured by antibody-focused scoring (Banerjee et al., 2017).

The domain I substitution E138K, widely discussed in the context of attenuation-associated phenotypes—similarly showed modest effects on E protein structure and antibody engagement in our in-silico assays. MD indicated overall RMSD behavior comparable to wild-type with only a small, local flexibility change near the mutation site, and docking/MM-GBSA did not reveal a substantial binding penalty (Kaiser et al., 2019). FoldX likewise suggested minimal impact on folding stability. This profile is consistent with a mutation that can alter viral fitness or virulence through mechanisms other than direct epitope disruption, while largely preserving the antigenic surface, which would be compatible with maintained immunogenicity in a vaccine background (Yang et al., 2017).

Considering together, the integrated MD–energy–stability evidence supports a practical hierarchy among the tested sites: EDIII mutations (S327Q, A366S) are the most likely to measurably shift neutralising-antibody interactions through localized increases in modest stabilization that reshape the epitope landscape, whereas T129M and E138K appear largely antigenically conservative in this framework. These findings identify candidate envelope substitutions whose structural context justifies further experimental characterization, including neutralization assays with polyclonal sera and population-level vaccine-effectiveness studies toward circulating genotypes and/or adopting multivalent designs, and they highlight EDIII-focused, structure-guided immunogen strategies (including epitope stabilization) as a targeted route to broaden protection without requiring wholesale changes to the overall E protein scaffold.

## Conclusion

A comprehensive comparative assessment of this study offers detailed mechanistic insight into the structural and immunological consequences of four key JEV E protein mutations: A366S, E138K, S327Q, and T129M. Among these, S327Q and A366S induce significant stabilization of domain III and attenuate antibody binding affinity, aligning with their established roles in genotype I/IV antigenic divergence and potential immune escape. In contrast, T129M and E138K exhibit minimal perturbation to epitope conformation, with E138K primarily contributing to attenuation through functional modulation of viral entry rather than antigenic remodelling. These results underscore the structural basis for the reduced efficacy of GIII-derived vaccines against heterologous genotypes and highlight the necessity of updating immunogen design, either through genotype-inclusive antigens or rational epitope engineering, to achieve broad-spectrum protection against evolving JEV strains. This integrative framework bridges computational biophysics and immunovirology, providing a rational foundation for next-generation vaccine optimization.

## DECLARATIONS

### Authors’ Contributions

HT conducted the research (Investigation), performed data analysis (Formal analysis), interpreted results, and prepared the initial draft (Writing – original draft). KPS conceptualized and designed the study (Conceptualization, Supervision), and contributed to manuscript review and editing (Writing – review and editing). RKP has technically done writing – review and editing. YSS and VR contributed to manuscript review and editing (Writing – review and editing). SSP and JH are also providing technical input. NK assisted with data analysis (Formal analysis). AP and BRG were involved in manuscript review and editing (Writing – review and editing). All authors read and approved the final manuscript.

## Acknowledgments

The authors wish to acknowledge the Spatial Epidemiological Laboratory at ICAR-NIVEDI and the Director of ICAR-NIVEDI for furnishing the requisite facilities to conduct this work. Additionally, the authors would like to recognise that the research was supported by the Indian Council of Agricultural Research, under the Department of Agricultural Research and Education, Government of India. The authors acknowledge the National Disease Modelling Consortium, Indian Institute of Technology Bombay, for research support. This study was funded by the Gates Foundation (Grant No. INV 044445)

